# Mitochondrial respiratory complexes, excitation-contraction coupling and mTOR signaling transcriptional programs are dysregulated in cardiometabolic HFpEF

**DOI:** 10.64898/2026.10.07.757359

**Authors:** Abraham Méndez-Fernández, Angélica Leticia Barraza-Arellano, Víctor Treviño, Gerardo García-Rivas

## Abstract

Heart failure is a leading cause of death and disability worldwide. Nearly, half of the cases correspond to HF with preserved ejection fraction (HFpEF), which is predicted to become the most prevalent form of HF in the forthcoming years. In this study, we performed a bulk RNA-seq assay to study the transcriptomic profiles of the hearts of HFpEF mice. HFpEF animals consisted in C57BL/6 mic that were fed a high-fat diet and exposed to L-NAME in the drinking water during five or eight weeks. Controls received a standard chow diet. Differential expression analysis yielded 1535 differentially expressed genes in HFpEF hearts relative to controls. 71 out of 78 DEGs contained in the GO-CC “Mitochondrial Inner Membrane (GO:0005743)” were found to be down-regulated, mostly affecting respiratory complexes I, III, IV and V in a time-dependent manner. Overrepresentation analyses also suggested that the mTOR signaling cascade is switched on, while “Diabetic Cardiomyopathy”, “Cardiac Muscle Contraction” and “Cytoskeleton in Muscle Cells” KEGG pathways were detected to be affected, probably pointing at excitation-contraction coupling impairment. In summary, the transcriptomic program of the HFpEF hearts consisted in a widespread downregulation of the expression of the subunits of mitochondrial respiratory complexes, upregulation of mTOR signaling and deregulation of the genes related to the excitation-contraction coupling.

## Introduction

Heart failure (HF) is the final common path of several cardiovascular diseases and currently stands as a leading cause of death and disability worldwide (Khan et al., 2024). About 2% of the adult population is affected by HF in the US (Siontis et al., 2022), and it is estimated that the global prevalence of HF is close to 64 million people (James et al., 2018). In particular, HF with preserved ejection fraction (HFpEF) is forecasted to drive HF prevalence in the near coming years due to the aging of the population and the rising incidence of obesity and diabetes (Dunlay et al., 2017). As obesity is virtually universal in the HFpEF patient (Packer, 2025), often co-occurring with hypertension (Packer et al., 2025), preclinical models combining both insults, metabolic plus hemodynamic stress, have been praised as reliable models for studying a HFpEF-like syndrome characterized by visceral adiposity, diastolic dysfunction, exercise intolerance and normal ejection fraction (Gao et al., 2024; Roh et al., 2022; Schiattarella et al., 2019).

Recent transcriptomic studies in mice models of HFpEF have unveiled significant alterations in cardiac gene expression. In a recently developed two-hit model that integrates diet-induced obesity with hypertension, achieved by renin overexpression, RNA sequencing revealed a transcriptomic signature characterized by activation of fibrotic pathways and downregulation of branched-chain amino acid catabolism (Berger et al., 2025). Comparative transcriptomic analyses across two HFpEF murine models, the high-fat diet (HFD) + L-NAME and the db/db + aldosterone mice, have identified significant etiology-and sex-specific differences in gene expression with, overall, little overlap between enriched pathways. However these results emphasize molecular heterogeneity of HFpEF, both models experienced downregulation of carbohydrate and upregulation of lipid metabolism (Hegyi et al., 2025). Collectively, the transcriptomic signature of HFpEF models seems to encompass profound remodeling of metabolism and contractile function, depending on HFpEF etiology. In the present study, we performed a bulk RNA-seq assay of ventricular tissue of HFD+L-NAME mice at two different stages across HFpEF progression, which may further expand our understanding of HFpEF molecular trajectory.

## Methods

### Animal model and treatment

CTRL animals consisted in eight to twelve-week-old male C57BL/6 mice fed with standard chow diet (PicoLab Rodent Diet 20 5053, LabDiet) and were given filtered water to drink. Conversely, HFpEF animals were fed with HFD (58Y1 - DIO Rodent Purified Diet w/60% Energy from Fat/Blue, TestDiet) and were simultaneously exposed to L-NAME (0.5 g/L) in their drinking water during five or eight weeks. Dietary and surgical procedures were approved by the institutional committee of animal care and use of the School of Medicine of Tecnológico de Monterrey (protocol 2021-005) and were executed in adherence to the Mexican regulation (NOM-062-ZOO-1999).

### RNA-Seq library preparation and sequencing

Upon the endpoints of either five or eight weeks of HFD + L-NAME exposure, total RNA was purified from about 30 mg of cryopulverized heart tissue using the TRIzol reagent (ThermoFisher Scientific). RNA was suspended in nuclease-free water and quantified fluorometrically (Qubit, ThermoFisher Scientific). RNA integrity was confirmed by sharp 28S and 18S rRNA bands after running a sample in a 1%-agarose gel. RNA-seq libraries were prepared following the manufacturer’s protocol with the TruSeq Stranded Total RNA Library Prep Kit (REF 20020737, Illumina) departing from 100 ng of total RNA. Each library corresponded to a single animal. Library preparation consisted of a first step of mRNA enrichment and fragmentation of the sample, followed by double-stranded cDNA synthesis and the ligation of adapters with unique identifiers for each library (TruSeq RNA Single Indexes Set A, REF 15032612, Illumina). Concentration of the double-stranded DNA libraries was measured fluorometrically (Qubit, ThermoFisher Scientific). Prior to sequencing, libraries were freshly diluted to a concentration of 4 nM and then pooled together in a centrifuge tube by taking 10 μL of each library. Next, pooled libraries were denatured by mixing 5 μL of the pool with 5 μL of freshly prepared NaOH 2 N followed by a thorough vortex and a five-minute incubation at room temperature.

Finally, 990 μL of the HT1 buffer was added resulting in 1 ml of 20 pM denatured libraries, of which 600 μL were loaded on a MiSeq Reagent Kit v3 cartridge (REF 15043893, Illumina). 150-bp paired-end reads were obtained after sequencing the libraries in a MiSeq system (Illumina). RNA-sequencing data have been deposited in the NCBI GEO public repository (accession number GSE349574).

### Bioinformatic analyses

Paired-end reads in the FASTQ files were aligned to the mouse genome assembly GRCm39 with the BWA (version 0.7.17-r1188) mem tool. Resulting SAM files were converted to BAM files and sorted by coordinates with the samtools version 1.11 view and sort functions, respectively. An index.bai file was generated for visual inspection of the aligned reads via the Integrative Genomics Viewer (Robinson et al., 2011). Aligned reads were then quantified with FeatureCounts v2.0.1 at the gene meta-feature level, resulting in a count matrix in which the *i, j* entry corresponded to the number of reads uniquely aligned to the i-th gene for the j-th library. Counts were normalized using DESeq2 median of ratios which accounted for sequencing depth and RNA composition, thus becoming suitable for inter-sample comparison. Differential expression analysis was conducted with the DESeq2 1.38.3 package (Love et al., 2014) using a design that contrasted CTRL vs. HFpEF libraries. Differentially expressed genes (DEGs) were defined as those having a Fold Change (FC) of at least 2 in either direction (up-or down-regulated) and a False Discovery Rate (FDR)<0.05. Overrepresentation (ORA) analysis was performed with the enrichR 3.4 package using the “GO Biological Process 2025”, “GO Cellular Component 2025”, “GO Molecular Function 2025” and “KEGG 2026” gene set libraries (Jawaid, 2025). Overrepresented terms were sorted by adjusted p-values. Only terms with an adjusted p-value below 0.05 were considered for further interpretation and plotting. Gene Ratio was calculated as the Gene Count over the total amount of genes annotated in a certain term. Visualization of the overrepresented terms and pathways was carried out with the ggplot2 and GOPlot 1.0.2 (Walter et al., 2015) in R version 4.2.2 (R Foundation for Statistical Computing).

### RNA-sequencing and bioinformatics

Total RNA was purified by the TRIzol method and was used as the input to prepare bulk RNA-seq libraries with the TruSeq Stranded Total RNA kit. Libraries were sequenced as 150-bp paired-end reads (MiSeq v3). Reads were aligned to the GRCm39 mouse genome (BWA-MEM), quantified with FeatureCounts, and analyzed for differential expression with DESeq2 (CTRL vs. HFpEF). DEGs were defined as those genes bearing an absolute fold change ≥ 2 and FDR < 0.05. Over-representation analysis used enrichR against GO and KEGG gene sets (adjusted p-vlaue < 0.05).

## Data visualization

Analyses and data visualization were performed in R 4.2.2.

## Results

### HFpEF animals displayed an increased body weight related to adiposity and dyslipidemia, elevated blood pressure and consequent diastolic dysfunction and myocyte hypertrophy

Phenotypic data of these animals has been thoroughly described by Méndez-Fernández and colleagues. Succinctly, relative to CTRL, HFpEF animals exhibited increased adiposity, dyslipidemia, elevated blood pressure, pulmonary congestion and overt HFpEF (Méndez-Fernández et al., 2026).

### Differential expression analysis (DEA) yielded 1535 genes differentially expressed between HFpEF and CTRL hearts

PCA analysis of the normalized counts matrix showed that HFpEF8week libraries located the furthest from the CTRLs in the PC space, whereas HFpEF5wk libraries laid somewhere between such groups (Figure 1A). As expected, sample-to-sample distance was low between HFpEF libraries, depicted as the darker 7×7 square in the up-left region of the heatmap shown in Figure 1B, whereas distance tended to be larger between HFpEF and CTRL samples. Only the library identified as HFpEF5wk_04 tended to be closer to library CTRL_02 than to any other HFpEF libraries, which may suggest it resembled closely the CTRL group, at least from a transcriptomic point of view. DEA yielded 1535 DEGs, of which 800 were overexpressed in HFpEF, while 735 were downregulated (pink dots in Figure 1C and rows in Figure 1D).

**Figure 1.**
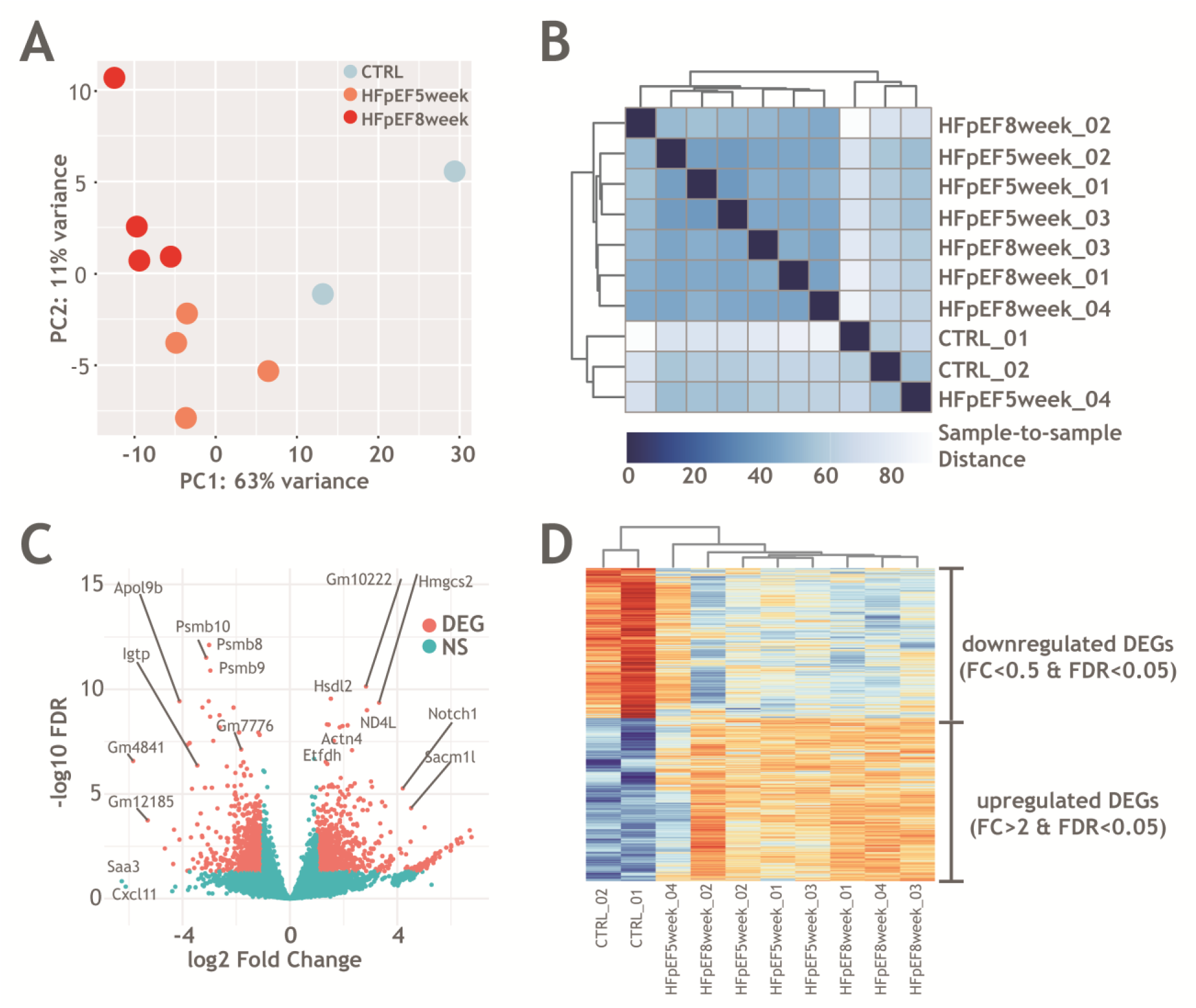
Differential-expression analysis (HFpEF vs. CTRL) yielded 1535 DEGs. **A.** Two-dimensional PCA projection of the normalized count matrix of the libraries in the RNA-seq dataset in PC1 and PC2. **B.** Sample-to-sample distance heatmap of the normalized count matrix of the libraries in the dataset. **C.** Volcano plot depicting DEGs (absolute FC≥2, FDR<0.05; CTRL as reference) as pink dots. **D.** Heatmap of gene expression as normalized counts scaled by row, in which each row is one DEG and each column is one RNA-seq library. DEGs are sorted by ascending FC.

### ORA analysis identified several terms and pathways related to mitochondrial metabolism, cytoskeleton, extracellular matrix organization, the TORC1/2 complexes and cardiac muscle contraction

Multiple GO terms related to mitochondria were unveiled by ORA analysis (Figure 2A), encompassing, Biological Process (BP) Cellular Respiration (GO:0045333), Mitochondrial ATP Synthesis Coupled Electron Transport (GO:0042775), Aerobic Electron Transport Chain (GO:0019646), Oxidative Phosphorylation (GO:0006119), Proton Motive Force-Driven Mitochondrial ATP Synthesis (GO:0042776), Aerobic Respiration (GO:0009060), Mitochondrial Electron Transport, NADH to Ubiquinone (GO:0006120), Proton Motive Force-Driven ATP Synthesis (GO:0015986), Energy Derivation by Oxidation of Organic Compounds (GO:0015980) and Mitochondrial Translation (GO:0032543) among the top 15 overrepresented terms based on the adjusted p-value. Cellular Component (CC) Mitochondrial Inner Membrane (GO:0005743) and Mitochondrial Membrane (GO:0031966) were the top 2 overrepresented terms based on adjusted p-value. No Molecular Function (MF) terms were overrepresented (adjusted p-vlaue>0.05). Other enriched terms were related to other important aspects of heart biology such as Collagen-Containing Extracellular Matrix (GO:0062023), TORC1 Complex (GO:0031931) and TORC2 Complex (GO:0031932) were also amid the top 15 CC overrepresented terms. Similarly, we conducted an ORA analysis using the KEGG 2026 database, which identified 24 overrepresented pathways encompassing Oxidative Phosphorylation, Diabetic Cardiomyopathy, Cardiac Muscle Contraction and Cytoskeleton in Muscle Cells (see Figure 2B). The full dataset derived from ORA analyses is available in Supplemental Table 1.

**Figure 2.**
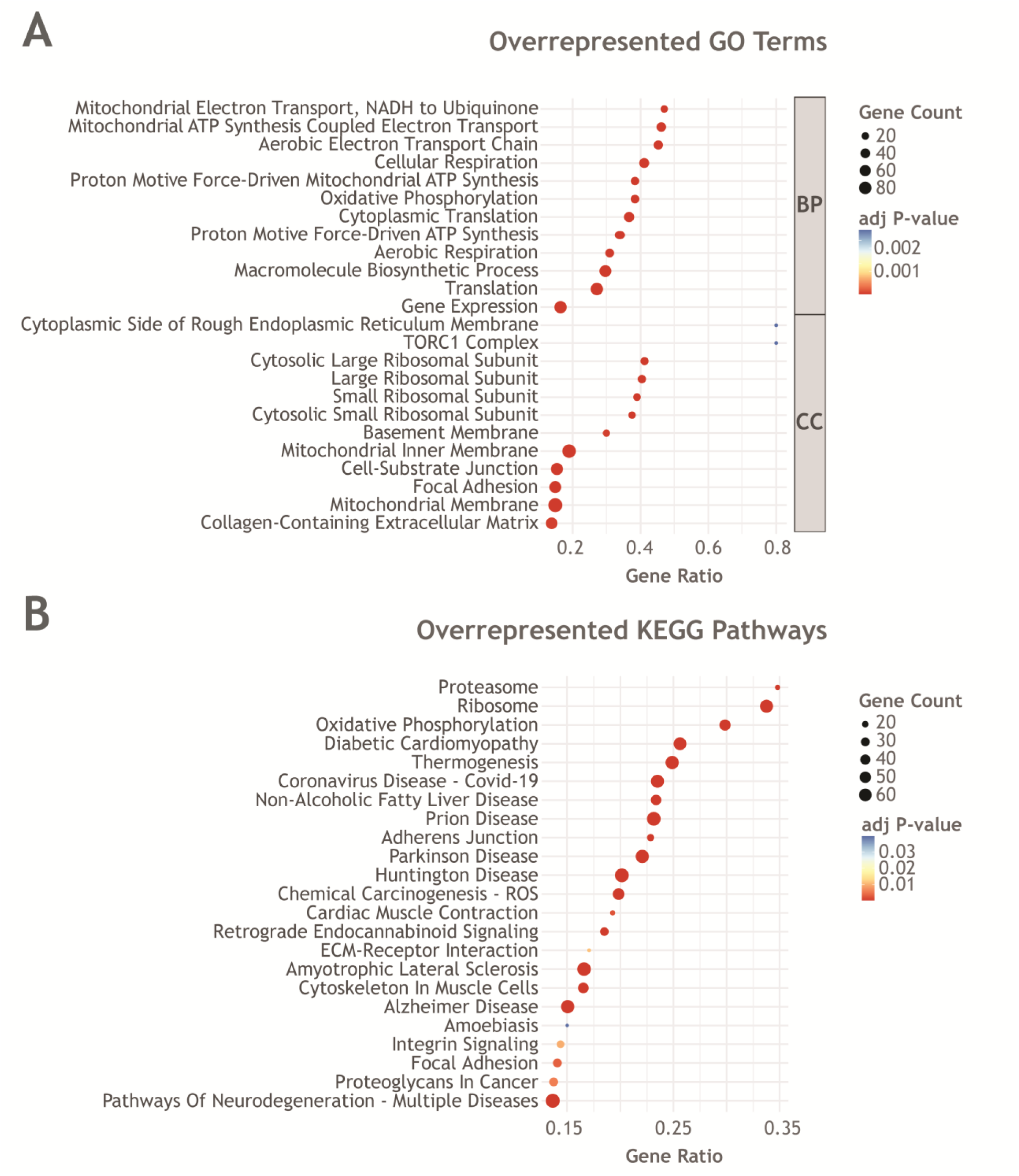
Enrichment analysis unveiled several mitochondria-related terms. **A.** Top 12 overrepresented GO BP and CC terms sorted by Gene Ratio. **B.** Overrepresented KEGG pathways sorted by Gene Ratio.

### DEGs that encode for inner mitochondrial membrane proteins were downregulated in HFpEF hearts in a manner dependent on time of exposure to HFD + L-NAME. Respiratory complexes I, III, IV and V resulted particularly affected

Since mitochondrial oxidative metabolism was common to the top overrepresented GO terms, we extracted the DEGs contained in the GO CC term “Mitochondrial Inner Membrane (GO:0005743)” to look in what direction the gene expression was actually altered. As shown in the heatmap of Figure 3A, the vast majority of DEGs, 71 out of 78 DEGs, contained in such a term, were downregulated. The chord plot in Figure 3B shows that DEGs encoding proteins of the inner mitochondrial membrane were indeed downregulated and that such genes were also annotated in the GO terms “Mitochondrial ATP Synthesis Coupled Electron Transport (GO:0042775)”, “Mitochondrial Electron Transport, NADH to Ubiquinone (GO:0006120)”, “Mitochondrial Electron Transport, Cytochrome C to Oxygen (GO:0006123)” and “Mitochondrial Electron Transport, Ubiquinol to Cytochrome C (GO:0006122)”. Thus, there is an overall downregulation of expression of the genes encoding for subunits of the respiratory complexes I, III, IV and V, which corresponding log2FC and FDR values can be found in Table 1. DEGs encoding for the subunits of the ATP synthase complex V exhibited the larger negative FC values. Interestingly, none of the four genes encoding for the succinate dehydrogenase complex II was differentially expressed.

**Figure 3.**
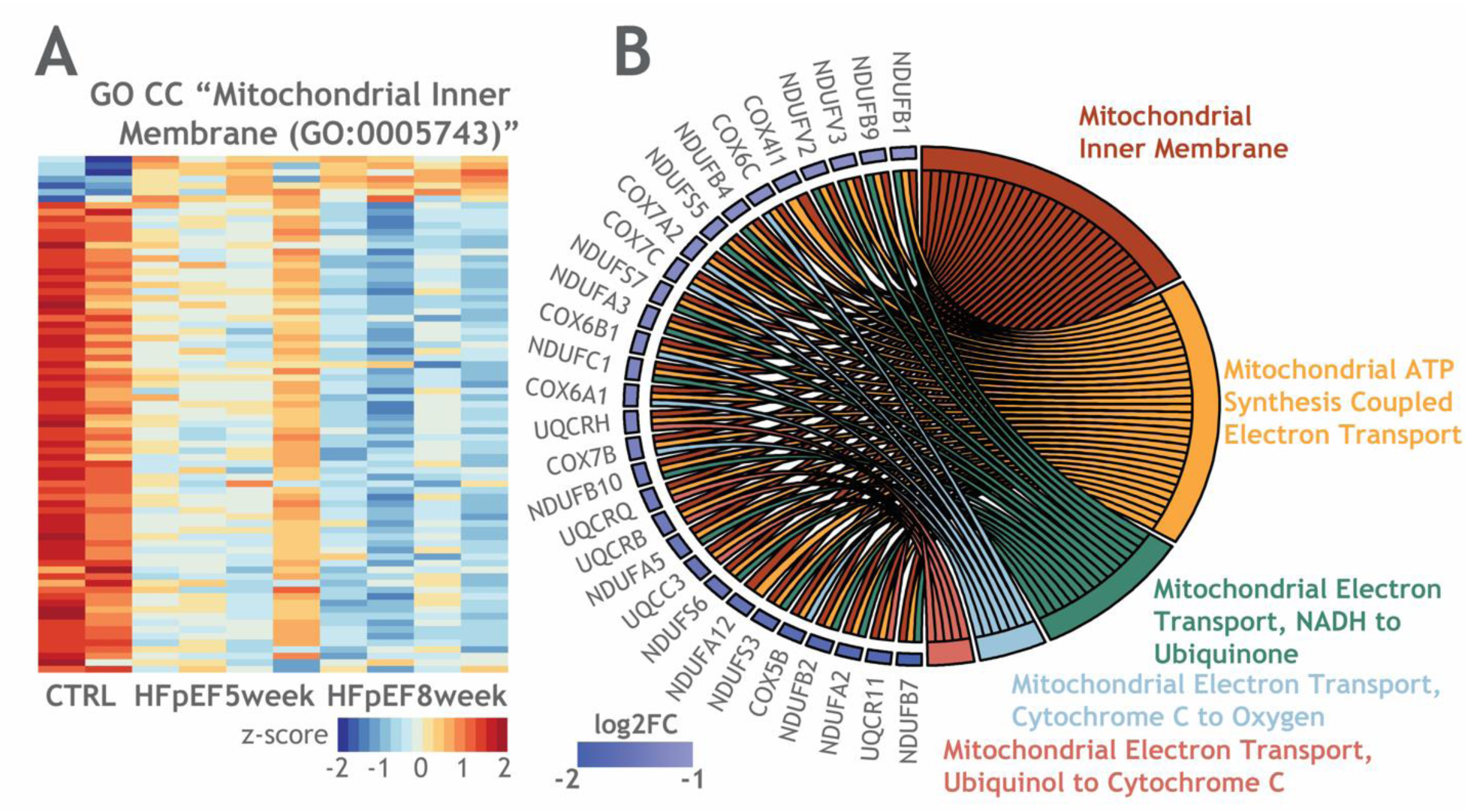
A. Heatmap of DEGs contained in the GO CC “Mitochondrial Inner Membrane (GO:0005743)” term showing 71 out of 78 DEGs were downregulated. Each column is one RNA-seq library and rows are DEGs sorted by ascending FC. **B.** Chord plot outlining the relationships of the DEGs contained in the “Mitochondrial Inner Membrane (GO:0005743)” GO CC term with the “Mitochondrial ATP Synthesis Coupled Electron Transport (GO:0042775)”, “Mitochondrial Electron Transport, NADH to Ubiquinone (GO:0006120)”, “Mitochondrial Electron Transport, Cytochrome C to Oxygen (GO:0006123)”, “Mitochondrial Electron Transport, Ubiquinol to Cytochrome C (GO:0006122)” GO terms.

**Table 1.**
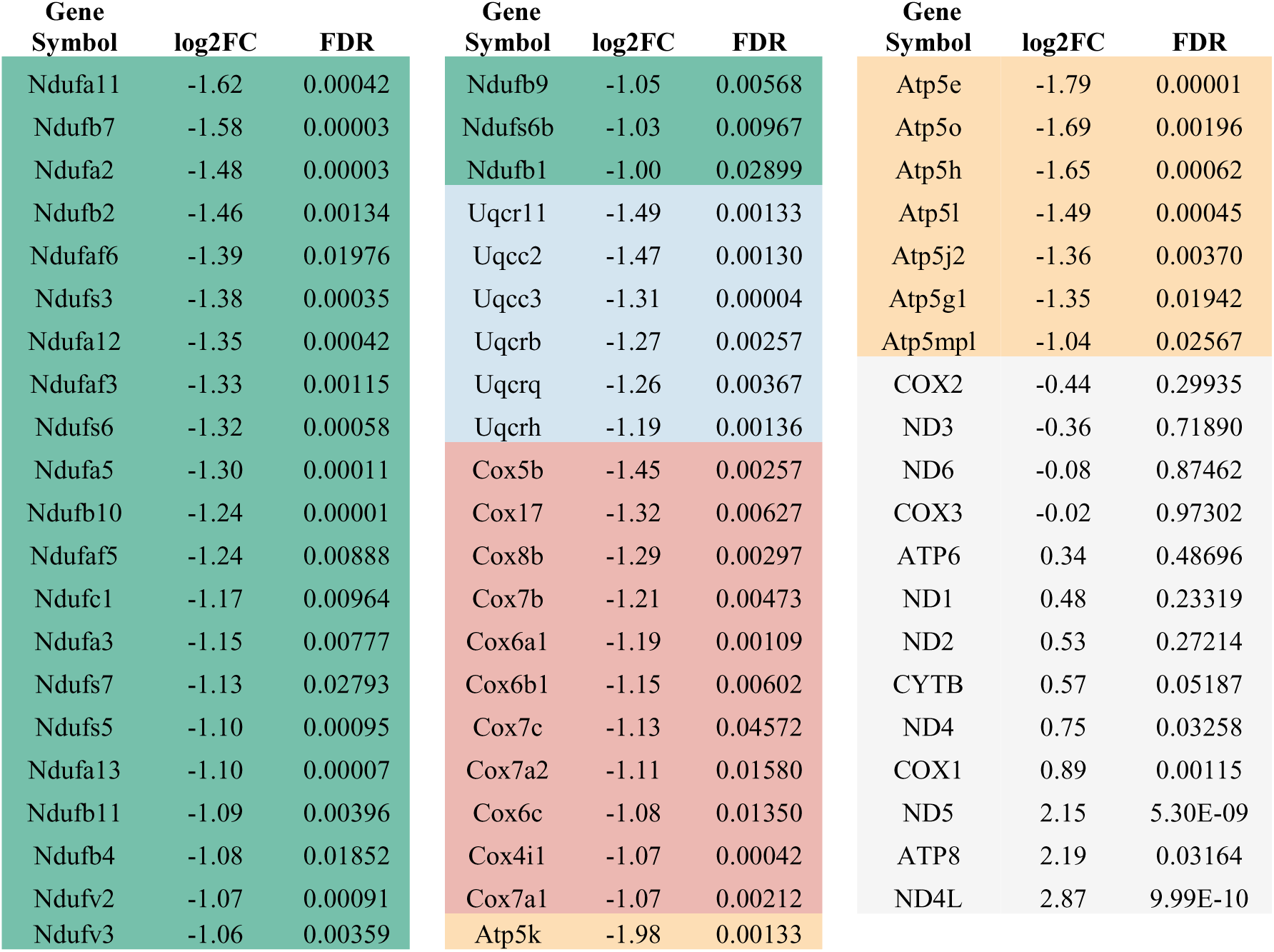
Nuclear-encoded DEGs affecting respiratory complexes I (green), III (light blue), IV (red), V (yellow) followed by the list of the 13 mt-DNA-encoded genes (grey) with their associated log2FC and FDR values. Each section is sorted by ascending FC relative to CTRL.

| Gene<br>Symbol | log2FC | FDR | Gene<br>Symbol | log2FC | FDR | Gene<br>Symbol | log2FC | FDR |
| --- | --- | --- | --- | --- | --- | --- | --- | --- |
| Ndufa11 | -1.62 | 0.00042 | Ndufb9 | -1.05 | 0.00568 | Atp5e | -1.79 | 0.00001 |
| Ndufb7 | -1.58 | 0.00003 | Ndufs6b | -1.03 | 0.00967 | Atp5o | -1.69 | 0.00196 |
| Ndufa2 | -1.48 | 0.00003 | Ndufb1 | -1.00 | 0.02899 | Atp5h | -1.65 | 0.00062 |
| Ndufb2 | -1.46 | 0.00134 | Uqcr11 | -1.49 | 0.00133 | Atp5l | -1.49 | 0.00045 |
| Ndufaf6 | -1.39 | 0.01976 | Uqcc2 | -1.47 | 0.00130 | Atp5j2 | -1.36 | 0.00370 |
| Ndufs3 | -1.38 | 0.00035 | Uqcc3 | -1.31 | 0.00004 | Atp5g1 | -1.35 | 0.01942 |
| Ndufa12 | -1.35 | 0.00042 | Uqcrb | -1.27 | 0.00257 | Atp5mpl | -1.04 | 0.02567 |
| Ndufaf3 | -1.33 | 0.00115 | Uqcrq | -1.26 | 0.00367 | COX2 | -0.44 | 0.29935 |
| Ndufs6 | -1.32 | 0.00058 | Uqcrh | -1.19 | 0.00136 | ND3 | -0.36 | 0.71890 |
| Ndufa5 | -1.30 | 0.00011 | Cox5b | -1.45 | 0.00257 | ND6 | -0.08 | 0.87462 |
| Ndufb10 | -1.24 | 0.00001 | Cox17 | -1.32 | 0.00627 | COX3 | -0.02 | 0.97302 |
| Ndufaf5 | -1.24 | 0.00888 | Cox8b | -1.29 | 0.00297 | ATP6 | 0.34 | 0.48696 |
| Ndufc1 | -1.17 | 0.00964 | Cox7b | -1.21 | 0.00473 | ND1 | 0.48 | 0.23319 |
| Ndufa3 | -1.15 | 0.00777 | Cox6a1 | -1.19 | 0.00109 | ND2 | 0.53 | 0.27214 |
| Ndufs7 | -1.13 | 0.02793 | Cox6b1 | -1.15 | 0.00602 | CYTB | 0.57 | 0.05187 |
| Ndufs5 | -1.10 | 0.00095 | Cox7c | -1.13 | 0.04572 | ND4 | 0.75 | 0.03258 |
| Ndufa13 | -1.10 | 0.00007 | Cox7a2 | -1.11 | 0.01580 | COX1 | 0.89 | 0.00115 |
| Ndufb11 | -1.09 | 0.00396 | Cox6c | -1.08 | 0.01350 | ND5 | 2.15 | 5.30E-09 |
| Ndufb4 | -1.08 | 0.01852 | Cox4i1 | -1.07 | 0.00042 | ATP8 | 2.19 | 0.03164 |
| Ndufv2 | -1.07 | 0.00091 | Cox7a1 | -1.07 | 0.00212 | ND4L | 2.87 | 9.99E-10 |
| Ndufv3 | -1.06 | 0.00359 | Atp5k | -1.98 | 0.00133 |  |  |  |

Intriguingly, down-regulation of the expression of such DEGs appeared to take place in a progressive fashion as a function of the time of exposure to HFD + L-NAME. Scatter plots in Figure 4 represent the normalized counts for the three DEGs of complexes I, III, IV and V that exhibited the higher absolute FC values. Relative to CTRL, expression of all these DEGs was diminished after a five-week exposure of HFD + L-NAME, and expression further declined for the 8-week HFpEF samples.

**Figure 4.**
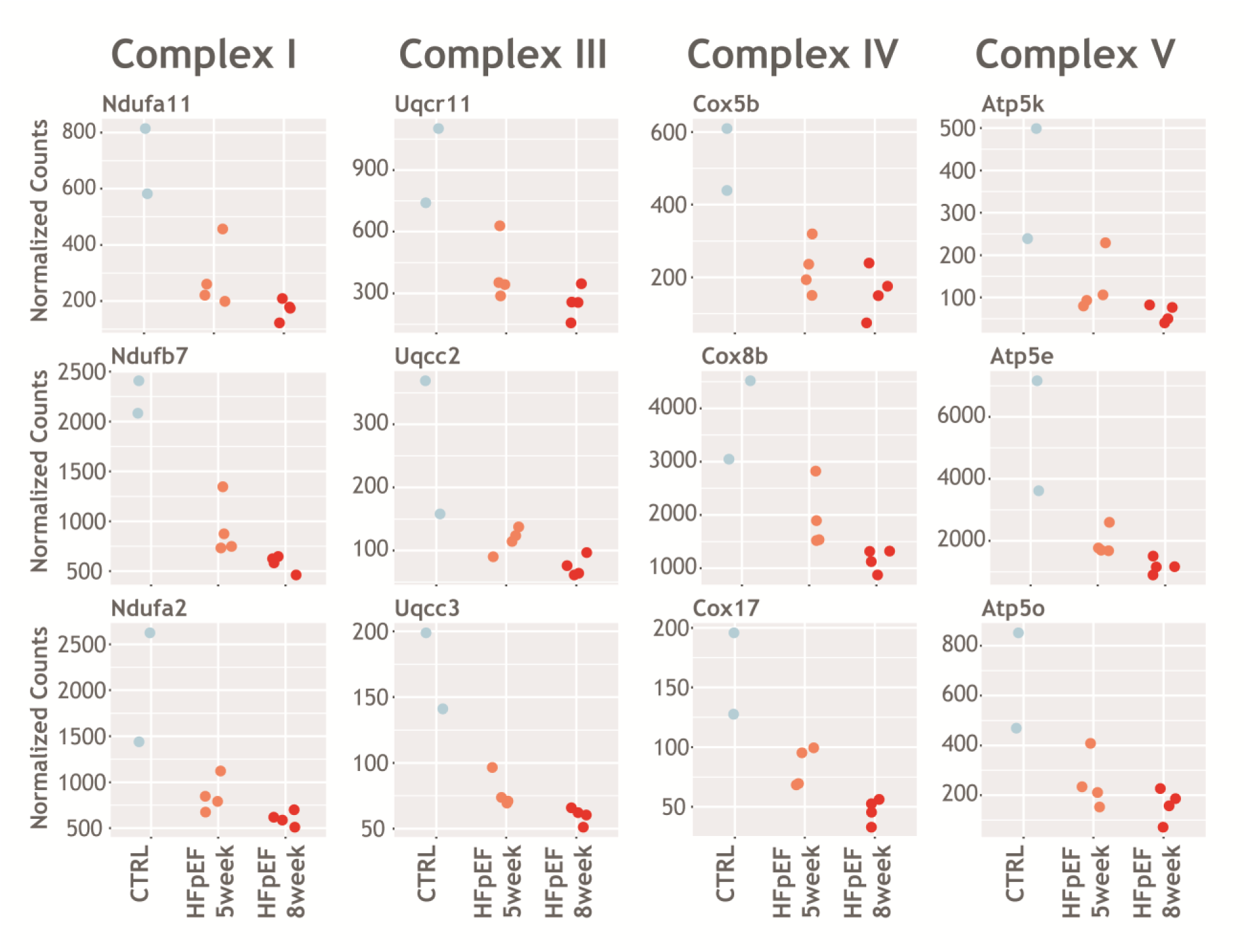
Expression of the DEGs encoding subunits of mitochondrial respiratory complexes I, III, IV and V (ATP synthase) declined in a fashion dependent of the time of exposure to HFD + L-NAME. Scatter plots of normalized counts resulting from the RNA-seq experiment, displaying the top three DEGs (based on FC) encoding for subunits of respiratory complexes I, III, IV and V. Blue dots represent CTRL animals, orange and red dots represent animals subjected to a HFD + L-NAME regimen during five (HFpEF 5 week) or eight weeks (HFpEF 8 week), respectively.

### The mTOR signaling pathway was found to be switched on

Both GO CC TORC1 (GO:0031931; adjusted p-value= 0.0028) and TORC2 (GO:0031932; adjusted p-value= 0.016) complexes were found to be significantly overrepresented. Indeed, several DEGs, encompassing Mtor itself, were significantly upregulated as summarized in Table 2. Interestingly, genes encoding the negative regulators of the mTOR pathway, such as Atm and Prkaa2, were also overexpressed, probably in a compensatory effort to limit excessive mTOR signaling.

**Table 2.** DEGs involved in the mTOR signaling pathway. DEGs were sorted by their corresponding FC value in a descendent fashion.

| **Gene**<br>**Symbol** | **log2FC** | **FDR** |
| --- | --- | --- |
| Atm | 6.24 | 0.00193 |
| Rps6ka5 | 3.35 | 0.03052 |
| Mtm1 | 2.26 | 0.00795 |
| Mlst8 | 2.19 | 0.04825 |
| Rps6kb1 | 1.99 | 0.00007 |
| Rictor | 1.90 | 0.03453 |
| Rps6ka4 | 1.70 | 0.01762 |
| Rps6ka2 | 1.54 | 0.03173 |
| Pdpk1 | 1.52 | 0.01333 |
| Usp9x | 1.51 | 0.00077 |
| Smg1 | 1.46 | 0.02429 |
| Fnip1 | 1.44 | 0.02767 |
| Sos2 | 1.37 | 0.00410 |
| Mtor | 1.23 | 0.00154 |
| Tti1 | 1.16 | 0.02148 |
| Rptor | 1.06 | 0.04070 |
| Prkaa2 | 1.02 | 0.00119 |
| Rps6 | -1.12 | 0.01305 |
| Lamtor2 | -1.21 | 0.01163 |
| Atp6v1g1 | -1.22 | 0.00456 |

### The excitation-contraction coupling process of the myocardium may be compromised in the HFpEF heart

Given that pathways specific to cardiac pathophysiology resulted overrepresented when exploring the KEGG database, namely, “Diabetic Cardiomyopathy”, “Cardiac Muscle Contraction” and “Cytoskeleton in Muscle Cells”, visualization of the network around these nodes was built (Figure 5). Most of the DEGs contained in “Diabetic Cardiomyopathy” encoded subunits of the respiratory complexes which, as mentioned above (Figures 3, 4), were mainly downregulated. Moreover, the CD36 fatty acid transporter, the angiotensin-converting enzyme (ACE) and the MTOR encoding genes were found to be differentially up-regulated. On the other hand, almost all the DEGs contained in “Cytoskeleton in Muscle Cells” were upregulated, which would be consistent with an ongoing hypertrophic program. Finally, while the vast majority of the DEGs in “Cardiac Muscle Contraction” overlapped with those of “Diabetic Cardiomyopathy”, it is worth noting that the expression of RYR2, TNNI3 and MYL3, key regulators of the calcium handling and sarcomere contraction, was significantly affected, suggesting anomalies in the myocardium excitation-contraction coupling process.

**Figure 5.**
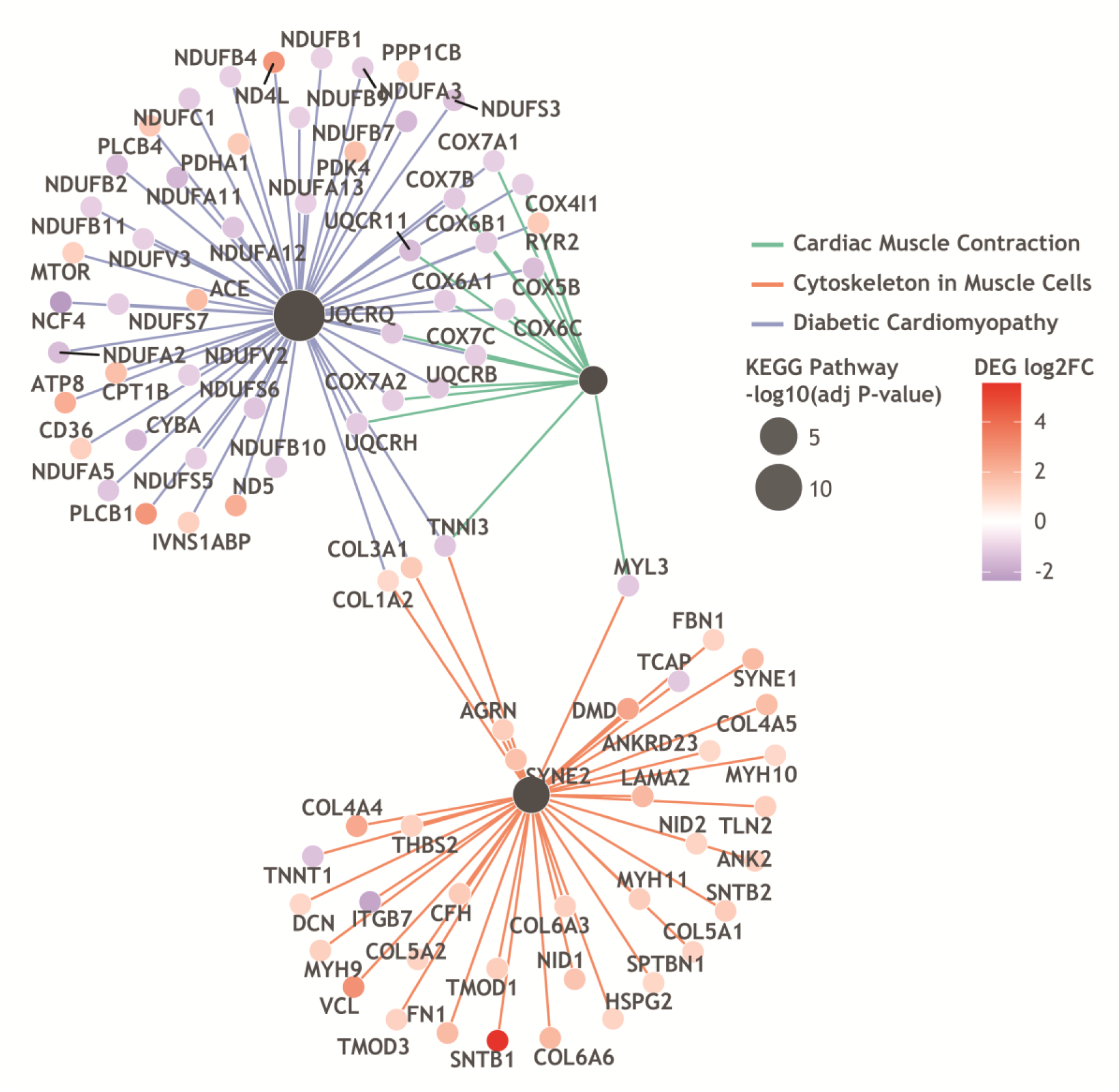
Gene network of heart-specific overrepresented KEGG pathways. Network in which nodes are the “Diabetic Cardiomyopathy”, “Cardiac Muscle Contraction” and “Cytoskeleton in Muscle Cells” KEGG pathways. The size of the node is the-log10 (adjusted p-value), while the FC value of the DEGs is color-coded.

## Discussion

### A widespread downregulation of genes encoding for the subunits of mitochondrial respiratory complexes is the transcriptomic signature of the cardiometabolic HFpEF heart

The most robust finding of the present study was a widespread down-regulation of the transcription of genes encoding for the mitochondrial respiratory complexes in the HFpEF heart. With the notable exception of complex II subunits, which expression was unaffected, numerous complex I, III, IV and V subunits experienced profound decreases in transcript levels (Table 1). Such a finding is consistent with a recent report, in which downregulation of mitochondrial enzymes involved in oxidative metabolism pathways was reported in the setting of HFpEF, particularly exacerbated in patients with severe obesity (Jani et al., 2025). The effect of obesity over the downregulation of genes encoding enzymes of the mitochondrial metabolism in HFpEF has been observed in the ZSF1 obese when compared against the ZSF1 lean rat (Summer et al., 2020), even when both ZSF1 obese and lean animals exhibited similar levels of hypertrophy and hypertension, when compared against Wistar Kyoto rats (Gibb et al., 2025). Particularly, in HFpEF murine preclinical models, dysregulation of the expression of genes implied in lipid and carbohydrate metabolism was found to be a common feature of the db/db plus aldosterone and the HFD + L-NAME mice (Hegyi et al., 2026). Hence, the present findings add to the growing amount of clinical and preclinical evidence pointing at a reconfiguration of gene expression in HFpEF that may ultimately affect the assembly and function of the respiratory complexes, ultimately manifesting as dysfunctional mitochondria, a well-known signature of the HFpEF heart (Schwartz et al., 2022).

Deviations from this notion have as well been reported, for instance in a feline model of progressive aortic constriction that experienced an early (one-month post-banding) decrease in mitochondrial gene expression (ND6, CHCHD3 and ATP5F1B only) that was followed by a shift from oxidative to glycolytic metabolism and normalization of the expression of the aforementioned genes at four-months post-banding (Gibb et al., 2021). However, in this case, a metabolic component of the HFpEF-like syndrome was completely absent and, therefore, results are hardly generalizable to HFpEF. Indeed, amid aortic stenosis patients, type-2 diabetes produced a distinctive transcriptomic signature of the left ventricle myocardium that comprised a downregulation of respiratory chain organization and function, relative to non-diabetics (Cherpaz et al., 2024). In humans, an early transcriptomics assay performed with right ventricle biopsies of patients characterized by obesity (median BMI was 41) and hypertension (98% of the patients were hypertense) pointed to the upregulation of the oxidative phosphorylation pathway in the HFpEF heart (Hahn et al., 2021), however a later high-throughput proteomics assay performed in the same samples showed directionality discordance between transcript and protein changes and concluded that protein abundance of mitochondrial proteins was indeed largely depressed in HFpEF relative to healthy controls (Jani et al., 2025).

Opposite to nuclear-encoded genes, genes encoded in the mitochondrial genome ND4L, ATP8, ND5, COX1 and ND4 (Table 1) were found to be differentially overexpressed (CYTB was at the borderline of significance, FDR=0.052). Similar observations were reported by Chen and colleagues in which despite diminished biogenesis, a possible compensatory upregulation in the transcription of mitochondrially-encoded genes was observed in HFD + L-NAME HFpEF mouse cardiomyocytes (Chen et al., 2022).

### The mTOR signaling pathway may be involved in the pro-hypertrophic response due to hemodynamic stress and nutrient excess

mTOR signaling cascade was found to be switched on, as can be seen in Table 2. Classically, the mTOR kinase assembles with regulatory proteins to form either the MTORC1 or the MTORC2 complex. Some proteins, as Mlst8 and Tti1 are common to both macromolecular complexes, whereas Raptor and Rictor are specific for MTORC1 and MTORC2, respectively. MTORC1 activation has been found to be essential for the hypertrophic response in both physiological and pathological conditions (Sciarretta et al., 2022). In the setting of pressure overload, it has been observed that MTORC1 activates S6K kinases, which, in turn, phosphorylate the S6 ribosomal protein component, and ultimately promote protein synthesis required for myocyte hypertrophy (Shioi et al., 2003; McMullen et al., 2004). Moreover, a similar MTORC1 activation has been observed in the context of increased nutrient availability, as in the case of mice fed with HFD, in both isolated cardiomyocytes and aortic tissue, presumably due to a depressed AMPK signaling (Reho et al., 2021; Sun et al., 2020). It is noteworthy that we found not only RPS6KB1, the gene encoding the S6K1 kinase, but also other members of the ribosomal S6 kinase family (RPS6KA2, RPS6KA4 and RPS6KA5) to be differentially overexpressed in the HFpEF heart (Table 2). Because of the aforementioned data, it could be speculated that the observed hypertrophy could rely on MTORC1 activation due to pressure overload combined with nutrient excess. However, as phosphorylation plays a major role in the MTORC1 regulation, it would be necessary to verify the phosphorylation state of the implicated proteins to bring support to this hypothesis. On the other hand, we also observed RICTOR overexpression, the gene encoding Rictor, the core protein of the MTORC2 complex, as well as USP9X, a deubiquitinase that promotes Rictor-mTOR assembly (Wrobel et al., 2020). During hemodynamic stress, MTORC2 is believed to exert cardioprotective pro-survival signaling that might attenuate functional impairment of the heart (Sciarretta et al., 2015; Shende et al., 2016; Simonson et al., 2017). Since overactivation of MTORC1 and MTORC2 in disease models appears to lead to detrimental and beneficial effects, respectively, it would be of major interest to determine which mTOR branch predominates in the current HFpEF model, in order to propose a feasible intervention to limit maladaptive remodeling.

### The cardiomyocyte excitation-contraction coupling may be dysregulated in the cardiometabolic HFpEF heart

As seen in Figure 5, genes encoding key proteins involved in the excitation-contraction coupling were found to be affected. Though a straightforward prediction of how this process may actually change in the ventricular cardiomyocyte cannot be derived from the present data, a downregulation of genes encoding mitochondrial proteins may be consistent with a defective calcium reuptake and relaxation, as has been reported before (Runte et al., 2017). Consistent with a recent report (Hegyi et al., 2025), we also found SCN4B transcript levels to be upregulated (log2FC=1.49, FDR=0.0086) in HFpEF, which may be behind the heightened late sodium current (Medeiros-Domingo et al., 2007) largely reported in the setting of HFpEF (Mustroph et al., 2022). As experimental evidence concerning calcium handling and sarcomere shortening after HFpEF + L-NAME exposure is often contradictory, depending on the species, strain and the time of exposure to the insults (Fenwick et al., 2024), functional experiments are warranted in order to shed light over excitation-contraction coupling in HFpEF.

## Limitations

We acknowledge that the transcriptomic approach of the current study is limited in the quest for understanding HFpEF nature, and that further experimental assays are needed to challenge the hypotheses derived from the present assay. However, we think that this study may have identified some cellular mechanisms within the HFpEF heart that may guide future studies.

## Conclusion

HFpEF hearts experience a widespread time-dependent repression of the transcription of genes encoding the subunits of mitochondrial respiratory complexes, with concomitant deregulation of the expression of genes related to the excitation-contraction coupling and mTOR signaling.

## Acknowledgements

We would like to thank the staff of the Core Lab Genomics of Tecnológico de Monterrey for the assistance in the sequencing assay. We would like to thank SECIHTI and the Institute for Obesity Research of Tecnológico de Monterrey.

